# Repair-Mediated Path Filtering of Mutation Load in Multilocus Homeostatic Populations

**DOI:** 10.64898/2026.10.11.758278

**Authors:** Ben Hardisty

**Affiliations:** School of Biological Sciences, University of Utah, 257 1400 E, Salt Lake City, 84112, Utah, USA

**Keywords:** mutation–selection balance, genetic load, homeostatic regulation, multilocus dynamics, DNA repair, structured populations

## Abstract

We study a nonlinear multilocus mutation–selection–repair system for an asexual population maintained at exact homeostasis by state-dependent reproductive feedback. Each of ***n*** identical biallelic damage loci mutates with probability ***U*** at reproduction and is repaired at rate ***ν***, with per-locus reproduction and mortality factors ***ζ***_***ρ***_ ***<* 1** and ***ζ***_***γ***_ ***>* 1**. Permutation symmetry and feedback through damage-class frequencies reduce the **2**^***n***^-state dynamics exactly to an **(*n* + 1)**-class ODE system on a forward-invariant simplex. Without repair, the nonlinear equilibrium is obtained in closed form throughout the branch on which the undamaged class persists. The microstate distribution factorizes as a Bernoulli product measure, the damage number is binomially distributed, and the branch terminates at ***U***_***c***_ **= 1 − *ζ***_***ρ***_***/ζ***_***γ***_, independently of ***n***. Under ***n* → ∞, *U* → 0**, and ***nU* → Θ**, the damage number converges to a Poisson distribution with mean **Θ*/*(*ζ***_***γ***_ **− *ζ***_***ρ***_). With repair, we prove existence and local asymptotic stability of an analytic rare-mutation equilibrium branch and derive a recursion for the factor ***E***_***k***_ suppressing the leading mutation load of class ***k***. Its source weights follow a conditional **Binomial(*k, ζ***_***ρ***_***/ζ***_***γ***_ ) law independent of ***n*** for every ***n* ≥ *k***. The resulting suppression lies between the product 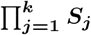, corresponding to sequential accumulation through every intermediate repair filter, and the terminal filter ***S***_***k***_ alone, corresponding to direct multilocus jumps that bypass them, where ***S***_***j***_ **= *λ***_***j***_***/*(*λ***_***j***_ **+ *jν*)** quantifies the balance between effective demographic depletion and repair. Repair also generates positive leading-order association among loci, breaking the no-repair product structure. Numerical integration of the full nonlinear system confirms the asymptotic predictions.

## 1 Introduction

Populations structured by genetic or molecular damage can occupy a large number of distinct states even when the underlying biological rules are simple. With *n* binary damage loci, the natural microstate description contains 2^*n*^ states, while homeostatic regulation can couple their frequencies nonlinearly through population-level demographic feedback. This raises two related mathematical questions: whether the resulting high-dimensional system admits an exact lower-dimensional description, and whether its equilibrium structure can be characterized analytically when mutation and repair act across multiple loci.

We address these questions in a haploid asexual population maintained at exact homeostasis by state-dependent reproductive feedback. Mutation occurs independently at undamaged loci during reproduction, while repair restores damaged loci in continuous time. Because the locus-level mutation and repair rules are identical and reproduction and mortality depend only on total damage, the full microstate vector field is equivariant under permutations of the loci, while its dynamics remain non-linear through the homeostatic feedback. As shown below, this structure nevertheless permits an exact reduction from the 2^*n*^ microstates to *n* + 1 damage classes and, in the absence of repair, an exact solution of the nonlinear equilibrium throughout the branch on which the undamaged class persists.

These questions arise naturally from mutation–selection balance. Haldane’s principle states that equilibrium genetic load is proportional to the deleterious mutation rate and, to leading order, independent of selection strength (Haldane 1937; Crow 1970). Subsequent deterministic theory extended Haldane-type load results to arbitrary mutation patterns among alleles and to quantitative genetic traits (Bürger and Hofbauer 1994), while haploid mutation–selection systems were analyzed for load and mutation-threshold phenomena (Hermisson et al. 2002). Under exact homeostatic regulation, however, the scale of the load is set by demographic turnover: in the absence of repair, the equilibrium load is the mutation probability per reproductive event multiplied by the realized per-capita turnover rate, giving *L*^∗^ = *εU* + *O*(*U* ^2^) for rare mutation, where *ε* = *γ*_0_*/ρ*_0_ (Hardisty 2026).

Repair introduces a second removal mechanism and changes this balance. In the one-dimensional damage hierarchy studied previously, repair from the first damaged class suppresses the leading equilibrium load by

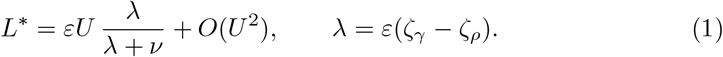

Thus *S*_1_ = *λ/*(*λ* + *ν*) quantifies the relative contribution of net demographic disadvantage and repair to the leading-order depletion of the first damaged class (Hardisty 2026). At leading order, repair reduces mutation load by adding a restorative loss term to the effective demographic depletion coefficient of the damaged class. This restoration process differs from reversible mutation in the genetic transmission process. Bürger and Hofbauer (1994) treat arbitrary mutation patterns, including back mutation, through a small-mutation expansion; in the classical discrete-allelic setting, selection-dependent corrections enter only at second order, while their multilocus quantitative-genetic treatment assumes global linkage equilibrium. Here, by contrast, *ν* is a continuous-time restoration rate acting on already-damaged loci, is held fixed as *U* → 0, and the no-repair product factorization is derived rather than assumed. This is also distinct from mutation-rate modifier models in which replication or repair machinery changes the underlying mutation rate (Sniegowski et al. 2000).

The preceding results are formulated on a one-dimensional damage hierarchy, in which mutation advances the population through ordered damage classes. They do not resolve damage across explicit loci, where a single reproductive event can create lesions at several previously undamaged sites. Many biological settings instead involve multiple loci accumulating damage in parallel — somatic mutations across tumor suppressor loci, epigenetic lesions at distinct regulatory sites, or molecular damage at multiple repair targets (Lindahl 1993; Chatterjee and Walker 2017; Hochedlinger and Jaenisch 2015; Knudson 1971). Multilocus mutation–selection balance has been studied in both asexual and recombining populations under a range of fitness architectures, including models organized by total mutation number and models with independent deleterious effects (Charlesworth 1990; Johnson 1999). Exact solutions for deterministic sequence-based mutation–selection models have also been obtained using statistical-mechanical methods (Baake and Wagner 2001). In that framework the usual mean-fitness normalization can be removed to yield a linear mutation–selection system, whereas the present dynamics are structurally different because the state-dependent feedback *R*(*f*) multiplies reproductive and mutational fluxes but not mortality or repair. A natural first step is therefore to determine whether explicit multilocus structure changes the no-repair equilibrium itself.

We analyze a haploid asexual population with *n* identical damage loci, multiplicative fitness across loci, per-locus mutation during reproduction, and independent per-locus repair, all held at exact homeostasis. Without repair the model is solvable in closed form throughout the pristine-positive branch, without a rare-mutation approximation. The equilibrium is an exact product measure: every locus is independently damaged with the same probability, the damage number is binomially distributed, and the exact load takes the same turnover form as in the one-step model, with the pristine-genome escape probability U = 1 − (1 − *U*)^*n*^ replacing the one-step mutation probability. The pristine-positive branch terminates at *U*_*c*_ = 1 − *ζ*_*ρ*_*/ζ*_*γ*_, the same threshold as in the single-locus model and independent of *n*. In the genomic rare-mutation limit *n* → ∞, *U* → 0 with *nU* → Θ, the binomial equilibrium converges to a Poisson distribution with mean Θ*/*(*ζ*_*γ*_ −*ζ*_*ρ*_), independent of the demographic ratio *ε*. This is the homeostatic counterpart of the classical result that multiplicative fitness yields a Poisson equilibrium distribution of deleterious mutations with mean equal to the mutation rate divided by the selection coefficient (Kimura and Maruyama 1966; Haigh 1978): exact homeostatic regulation leaves that form intact, sharpens it to an exact binomial at finite *n*, and replaces the selection coefficient by the per-locus reproduction–mortality contrast *ζ*_*γ*_ − *ζ*_*ρ*_.

With repair, we first establish the existence of an analytic rare-mutation equilibrium branch and show that this branch is locally asymptotically stable for sufficiently small *U* . For *n* ≥ 2 and *ν >* 0, the no-repair product factorization is not preserved on the rare-mutation branch for sufficiently small positive *U* . Although repair acts independently at each damaged locus, its class-level flux is *kν*, and its interaction with the geometrically varying reproductive and mortality rates prevents the no-repair binomial balance from carrying over unchanged. The resulting equilibrium therefore contains genuinely multilocus corrections, whose structure is revealed most clearly in the rare-mutation expansion below.

The main result quantifies this coupling. The leading load contribution from damage class *k* is suppressed relative to its no-repair value by a factor *E*_*k*_ obeying a weighted-average recursion over the source classes that feed class *k*. The single-locus result *E*_1_ = *S*_1_ is recovered exactly. For *k* ≥ 2, a purely sequential route 0 → 1→ · · · → *k* passes through every intermediate damage class and is filtered repeatedly, whereas simultaneous mutation of several loci in one reproductive event permits direct jumps *j* → *k* that bypass intermediate classes and hence bypass some filters. The realized suppression is therefore bounded between the fully sequential cascade product and the terminal filter alone:

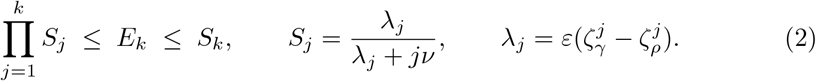

The mixture of routes admits a closed description. At leading order in *U*, the no-repair flux into class *k* is distributed over its source classes *j < k* according to a Binomial(*k, p*) law conditioned on *j < k*, with

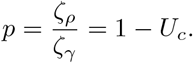

For every *n* ≥ *k*, this conditional route law is independent of the total number of loci. As the per-locus reproduction–mortality contrast strengthens (*p* → 0), the source weight concentrates on direct 0 → *k* multilocus jumps and *E*_*k*_*/S*_*k*_ → 1. As the contrast weakens (*p* → 1), the conditional source law instead concentrates on *j* = *k* − 1, corresponding to sequential accumulation through the immediately preceding class. In this latter limit the class-specific repair filters also vanish, so source-law concentration should not be interpreted as relative convergence of *E*_*k*_ to the full cascade product. Thus, at leading order, the number of loci sets how abundant each damage class is, but not the mixture of routes by which a fixed class *k* is reached, and therefore not its suppression factor *E*_*k*_. Repair also breaks the no-repair product structure in a definite direction: for distinct loci, the leading pairwise covariance on the rare-mutation branch is positive, so the repaired damage-number distribution is overdispersed relative to a binomial distribution with the same marginal damage probability.

These results also have a natural interpretation in multi-hit models of carcinogenesis, where repair depletes intermediate mutational states while simultaneous multilocus events can bypass them (Armitage and Doll 1954; Knudson 1971; Beerenwinkel et al. 2007; Foo et al. 2014; Portier and Kopp-Schneider 1991).

The paper is organized as follows. Section 2 introduces the *n*-locus model and reduces the microstate dynamics to a closed system on total damage classes. Section 3 solves the no-repair system exactly throughout the pristine-positive branch, without a rare-mutation approximation, and establishes the independence, binomial, and threshold results. Section 4 establishes the analytic rare-mutation equilibrium branch and its local stability, proves the path-filtering suppression theorem and its *n*-independent route law, derives the repair-induced association among loci, and compares the leading-order predictions with direct numerical integration of the full nonlinear system. Section 5 discusses biological implications, limiting cases, and extensions.

## 2 Multilocus homeostatic model

We extend the homeostatic mutation–selection framework developed in Hardisty (2026) by resolving damage across *n* explicit loci. The ingredients needed for the multi-locus analysis are restated in full below, and the results of Sections 3 and 4 are derived from them directly. The demographic interpretation of the feedback and mutation load is developed at greater length in the parent model, to which we refer for motivation rather than for any result required here.

### 2.1 State space and symmetry reduction

Consider a haploid asexual population carrying *n* biallelic loci. Each locus is either undamaged (0) or damaged (1), so a microstate is

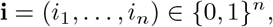

with total damage

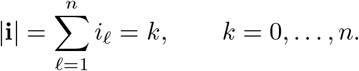

Let *g*_**i**_(*τ*) denote the frequency of microstate **i**, and define the aggregate damage-class frequencies

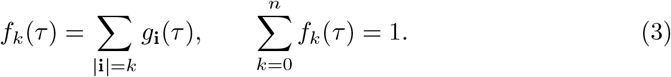

All loci are assumed identical with respect to mutation, repair, reproduction, and mortality. Reproduction and mortality therefore depend on a microstate only through its total damage:

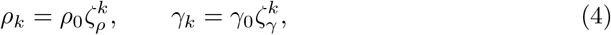

where

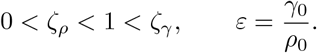

Because the transition rules are invariant under permutations of the loci, the microstate vector field is equivariant under the action of the symmetric group S_*n*_. Moreover, mutation, repair, reproduction, and mortality depend on a microstate only through its Hamming weight. The homeostatic feedback introduced below also depends only on the damage-class frequencies, because its numerator and denominator satisfy

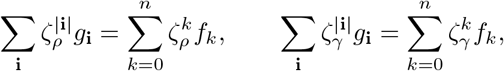

and therefore 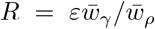 is determined entirely by (*f*_0_, …, *f*_*n*_). Thus any two microstates with the same Hamming weight have identical per-capita transition rates into each damage class, with a common feedback factor determined by the class frequencies. Summing the microstate equations over Hamming-weight classes therefore yields a closed system for *f*_0_, …, *f*_*n*_, without requiring exchangeable initial conditions. Exchangeable initial data additionally imply equal frequencies among microstates within each damage class, but this stronger condition is not required for the class-level dynamics below.

### 2.2 Homeostasis, mutation, and repair

Time is rescaled by the pristine reproduction rate, *τ* = *ρ*_0_*t*. Define the population-mean relative reproduction and mortality rates

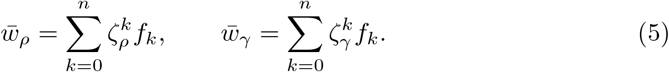

Exact homeostasis is imposed through the state-dependent reproductive feedback

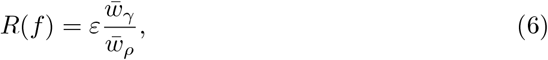

so that total reproductive and mortality fluxes are equal.

Mutation occurs during reproduction. Each undamaged locus mutates independently with probability *U* per reproductive event, with no reverse transitions in the offspring-mutation kernel. Thus a parent in damage class *j* produces an offspring in class *k* ≥ *j* with probability

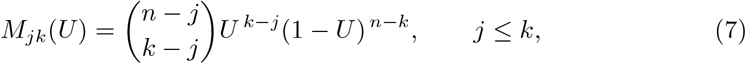

and *M*_*jk*_ = 0 for *k < j*. For each 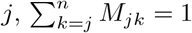.

Repair acts independently at each damaged locus at rate *ν* per unit dimensionless time. An individual in class *k* therefore moves to class *k ™* 1 at total rate *kν*. Repair is conservative: it redistributes individuals among damage classes but does not directly alter total population size. Throughout, *n* ≥1, 0 ≤*U* ≤1, *ν* ≥0, and *ρ*_0_, *γ*_0_ *>* 0, so *ε >* 0.

#### Interpretation of the damaged state

The binary state at each locus represents an effective deleterious damage state, rather than necessarily a permanently fixed nucleotide substitution. The parameter *U* specifies the probability that an undamaged locus enters this state in association with reproduction, while *ν* specifies its continuous-time restoration rate. Thus, repair describes reversal of a repairable lesion or other reversible molecular state within an individual, not the routine reversal of an established sequence mutation. The terminology of mutation–selection balance is retained because the upward transitions occur during reproduction and the resulting damage states have class-dependent demographic consequences. The model does not resolve the molecular steps separating initial DNA damage, mutation fixation, and restoration.

### 2.3 Closed class dynamics and mutation load

With the convention *f*_*n*+1_ = 0, the complete class-frequency system can be written compactly as

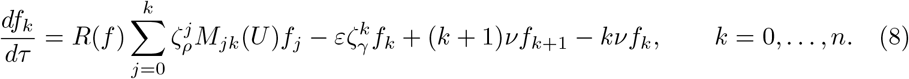

Let

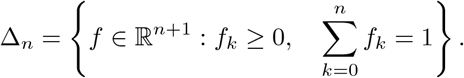

Summing (8) over *k*, using 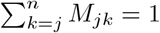, gives

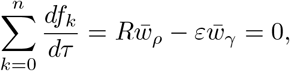

while the repair terms telescope. Thus the normalization ∑_*k*_*f*_*k*_ = 1 is preserved.

Nonnegativity is preserved as well. On any boundary face *f*_*k*_ = 0,

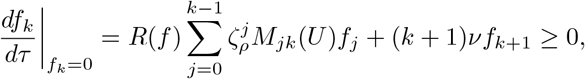

with the sum understood to be empty when *k* = 0. Hence the vector field is quasipositive on Δ_*n*_. Moreover,

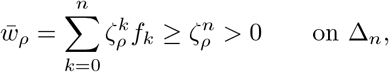

so the feedback *R*(*f*) is nonsingular there. Consequently, Δ_*n*_ is forward invariant, and for every initial condition *f* (0) ∈ Δ_*n*_ the system admits a unique global forward solution.

Structurally, Eq. (8) is a nonlinear ODE system on the standard *n*-simplex, with the nonlinearity entering through the single scalar feedback *R*(*f*). This feedback rescales reproductive and mutational fluxes, while mortality and repair remain linear. The system is therefore not generated by a fixed linear mutation–selection operator followed only by scalar mean-fitness normalization.

Following Hardisty (2026), define the fixed-reference mutation load by

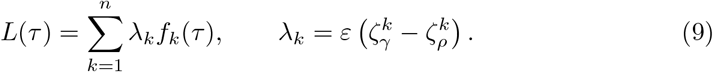

The load *L* is a fixed-reference demographic deficit, measured relative to pristine reproduction and mortality, rather than a normalized fraction of mean population fitness. The exact demographic identity derived in the parent model carries over unchanged:

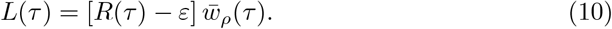

Indeed, 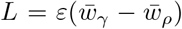 and 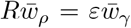 by (6). Thus the same load identity holds here despite the possibility of simultaneous mutation at several loci.

## 3 Exact no-repair equilibrium at finite mutation pressure

Before introducing repair, we solve the multilocus system exactly throughout the pristine-positive branch, without a rare-mutation approximation. Set *ν* = 0 and define

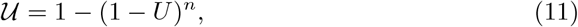

the probability that reproduction by a pristine individual produces at least one newly damaged locus.

### Proposition 1

(Exact no-repair multilocus equilibrium) *Set ν* = 0. *An equilibrium with* 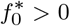*exists if and only if*

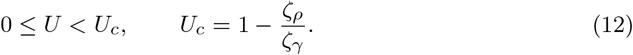

*When it exists, this equilibrium is unique*.

*On this branch, define*

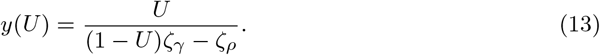

*Then*

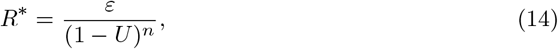

*and the equilibrium damage-class distribution is*

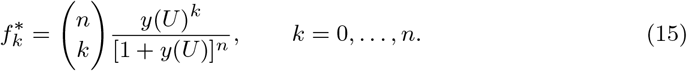

*The equilibrium load satisfies the exact identities*

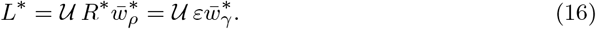

*Here* U *retains its interpretation as the pristine-genome escape probability*.^1^ *As U* ↑ *U*_*c*_, *y*(*U*) → ∞ *and* 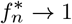.

*Proof* The class-0 equilibrium equation is

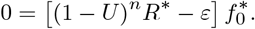

Since 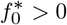,

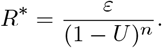

For 1 ≤ *k* ≤ *n*, the no-repair equilibrium equation can therefore be written as

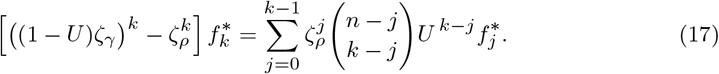

Now let

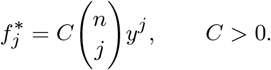

Using

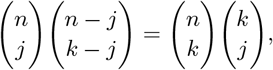

the right-hand side of (17) becomes

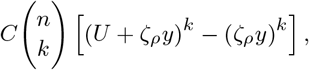

whereas the left-hand side becomes

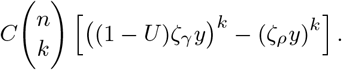

The two expressions agree for every *k* provided

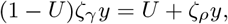

which gives (13). Normalization then yields *C* = (1 + *y*)^−*n*^ and hence (15). It remains to verify that the constructed distribution is consistent with the homeostatic feedback, since *R* is determined by *f* rather than being an independent parameter. From (15),

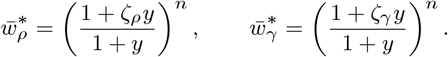

Hence

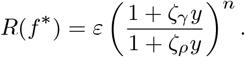

By (13),

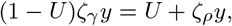

and therefore

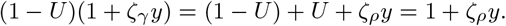

Thus

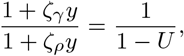

so

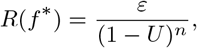

which agrees with the value obtained from the class-0 balance. Therefore the constructed distribution is a genuine equilibrium of the full nonlinear system.

For 0< *U* < *U*_*c*_

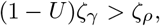

so *y >* 0 and all equilibrium frequencies are positive. Moreover, the coefficient multiplying 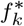 in (17) is positive for every *k*. The triangular system therefore determines each 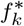uniquely as a positive multiple of 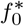, and normalization fixes 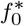 uniquely. At *U* = 0 the solution reduces to the pristine state.

If *U* = 1, the class-0 equilibrium equation reduces to 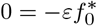, contradicting 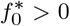. For *U*_*c*_≤ *U <* 1, the coefficient in the *k* = 1 equation is nonpositive, whereas its right-hand side is 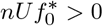. Hence no biologically admissible equilibrium with 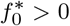 exists for *U* ≥ *U*_*c*_.

Finally,

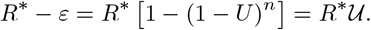

Substitution into the exact load identity (10), followed by 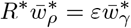, gives (16). Since the denominator of (13) tends to zero from above as *U* ↑ *U*_*c*_, *y* → ∞ and (15) gives 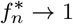. $#x25A1;

### Corollary 1

(Genomic mutation-escape threshold) *Under the assumptions of Proposition 1, the threshold U*_*c*_*in* (12) *is independent of n, whereas the corresponding critical probability that reproduction by a pristine individual produces at least one newly damaged locus*,

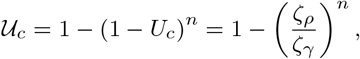

*increases with n and converges to one as n* → ∞. *This is precisely the branch endpoint* 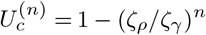 *of the one-dimensional n-class damage hierarchy (*Hardisty 2026*): expressed as pristine-genome escape probabilities, the multilocus genome and the capped damage hierarchy terminate at the same point*.

### Corollary 2

(Exact product-measure equilibrium) *Under the assumptions of Proposition 1, let*

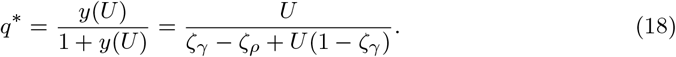

*Then the equilibrium distribution on the full microstate space* {0, 1}^*n*^ *is*

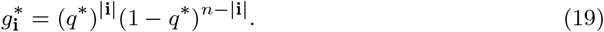

*Thus the n loci are independent and identically distributed Bernoulli*(*q*^∗^) *variables at equilibrium. In particular, the total damage number is exactly*

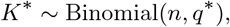

*and summing* (19) *over all microstates with* |**i**| = *k recovers* (15).

*Proof* Identify a microstate with its set *A* ⊆ {1, …, *n*} of damaged loci, and write 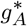 for its equilibrium frequency. If *B* is a target microstate with |*B*| = *k*, then in the absence of repair it can receive offspring only from states *A* ⊆ *B*. Its equilibrium balance is

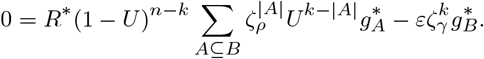

Using *R*^∗^ = *ε*(1 − *U*)^−*n*^ from Proposition 1 and separating the term *A* = *B* gives

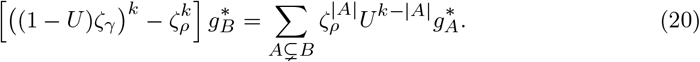

We prove by induction on *k* that

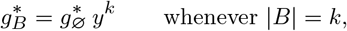

where *y* = *y*(*U*) is given by (13). The result is trivial for *k* = 0. Assuming it holds for all proper subsets of *B*, the right-hand side of (20) becomes

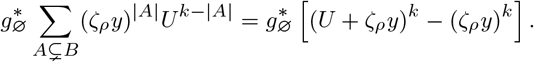

Since (13) implies

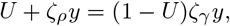

this is

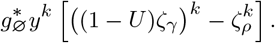

For *U < U*_*c*_the bracket is positive, so 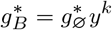 .

Normalization over all 2^*n*^ microstates gives

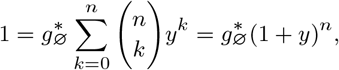

and hence

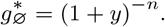

Therefore

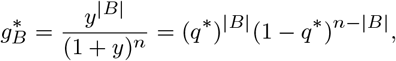

which is the product measure (19). Because these microstate frequencies aggregate exactly to (15), their population-mean reproduction and mortality rates, and hence the homeostatic feedback *R*, coincide with those already verified in Proposition 1. Thus no additional feedback-consistency condition is required at the microstate level. $#x25A1;

### Corollary 3

(Poisson limit) *Let n* → ∞ *and U* → 0 *with nU* → Θ *<* ∞. *Then, on the no-repair branch*,

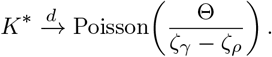

*In particular, the limiting mean is independent of ε*.

*Proof* From (18),

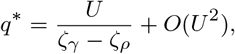

so

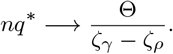

Since *K*^∗^ ∼ Binomial(*n, q*^∗^) by Corollary 2, the standard binomial–Poisson limit gives the result. □

### Compatibility with the genomic threshold

The scaling *nU* → Θ fixes the expected number of newly damaged loci per reproduction by a pristine individual, whereas the probability of at least one such event satisfies

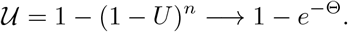

These quantities are distinct, as is the limiting equilibrium mean Θ*/*(*ζ*_*γ*_ − *ζ*_*ρ*_) in Corollary 3. Since 1 − *e*^−Θ^ *<* 1 whereas _*c*_ 1 by Corollary 1, the Poisson limit remains within the pristine-positive equilibrium regime for every fixed finite Θ and all sufficiently large *n*.

#### Proposition 2

(Complete no-repair class-equilibrium classification) *Set ν* = 0 *and* 0 ≤ *U* ≤ 1. *For each k* = 0, …, *n, the face*

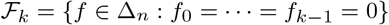

*is forward invariant*.

*If* 0 ≤ *U < U*_*c*_, *the class-frequency system has exactly n*+1 *equilibria f* ^[*k*]^, *one with lowest occupied class k for each k* = 0, …, *n. Writing*

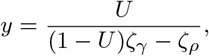

*these equilibria are*

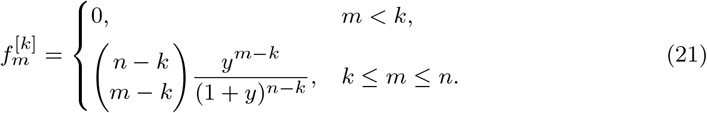

*Their homeostatic feedback values are*

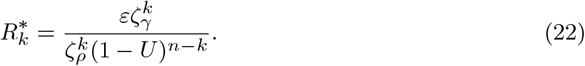

*For* 0 *< U < U*_*c*_, *the support of f* ^[*k*]^ *is exactly* {*k*, …, *n*}. *At U* = 0, *these equilibria reduce to the vertices e*_*k*_, *which are the only equilibria*.

*The case k* = 0 *recovers Proposition 1, while f* ^[*n*]^ = *e*_*n*_.

*For k* ≥ 1, *the eigenvalues transverse to* F_*k*_*at f* ^[*k*]^ *are*

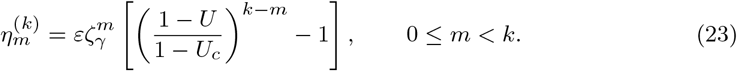

*They are strictly positive for U < U*_*c*_, *so every boundary equilibrium is unstable to perturbations introducing lower-damage classes*.

*For U*_*c*_≤ *U* ≤ 1, *the fully damaged state e*_*n*_*is the unique equilibrium in* Δ_*n*_. *Its eigenvalues are given by* (23) *with k* = *n. Consequently, e*_*n*_*is locally asymptotically stable for U > U*_*c*_*and nonhyperbolic at U* = *U*_*c*_, *where all its eigenvalues vanish*.

*Proof* With *ν* = 0, mutation never decreases damage class, so every F_*k*_is forward invariant.

Fix *k < n* and restrict the dynamics to F_*k*_. Set *d* = *n* − *k* and *g*_*s*_= *f*_*k*+*s*_for *s* = 0, …, *d*. Define the rescaled feedback

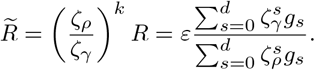

The restricted system is therefore the no-repair *d*-locus system under the positive time rescaling 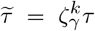. Since this time change leaves equilibria unchanged, Proposition 1 applies directly.

Proposition 1 therefore gives the unique equilibrium with *g*_0_ *>* 0 for *U < U*_*c*_. Its binomial formula yields (21), and the corresponding feedback yields (22). For *k* = *n*, the face consists only of *e*_*n*_, which is an equilibrium for every *U*.

To exclude additional equilibria, let *k* be the lowest occupied class of any equilibrium. For *U <* 1 and *k < n*, the class-*k* balance forces

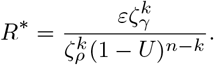

If *U*_*c*_≤ *U <* 1, the coefficient of 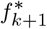in its equilibrium equation is

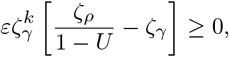

whereas mutation from class *k* supplies a strictly positive inflow. This is impossible. At *U* = 1, the balance of any lowest occupied class *k < n* gives 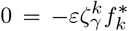 again a contradiction. Hence only *e*_*n*_remains for *U* ≥ *U*_*c*_.

At *U* = 0, no mutation occurs, so the equilibrium balance for each occupied class *m* requires

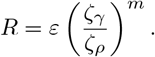

Since *ζ*_*γ*_*/ζ*_*ρ*_*>* 1, this condition cannot hold for two distinct occupied classes. Thus every equilibrium at *U* = 0 is a vertex *e*_*k*_.

Finally, linearize at *f* ^[*k*]^. For every *m < k*, the equilibrium has 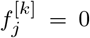for all *j* ≤ *m*. Consequently, derivatives of the feedback contribute nothing to the class-*m* equation at first order. The linearized equations for classes *m < k* form a lower-triangular block, whose diagonal entries are

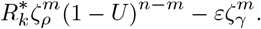

Substituting (22) and using *ζ*_*ρ*_*/ζ*_*γ*_= 1 − *U*_*c*_gives (23). The same calculation applies at *e*_*n*_for every *U*, including *U* = 1.

For *U < U*_*c*_, these eigenvalues are positive whenever *m < k*. For *U > U*_*c*_, all eigenvalues at *e*_*n*_are negative, making it a hyperbolic sink on Δ_*n*_. At *U* = *U*_*c*_, they all vanish. □

For *U < U*_*c*_, Proposition 2 shows that every equilibrium other than the pristine-positive branch is transversely unstable; global attraction of that branch is not claimed. For *U > U*_*c*_, the fully damaged state *e*_*n*_ is the unique equilibrium and is locally asymptotically stable.

## 4 Repair-mediated path filtering

We now restore repair and examine the equilibrium branch for rare mutation. The exact no-repair solution of Section 3 provides the baseline against which repair suppression can be measured.

### 4.1 Rare-mutation branch with repair

At *U* = 0, the pristine state

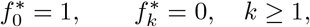

is an equilibrium for every finite repair rate *ν* ≥ 0. Recall from (9) that

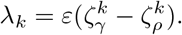

#### Proposition 3

(Rare-mutation hierarchy with repair) *For fixed n <* ∞ *and finite ν* ≥ 0, *there is a locally unique equilibrium branch analytic in U near U* = 0. *Along this branch*,

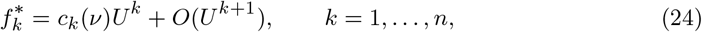

*with c*_0_(*ν*) = 1 *and*

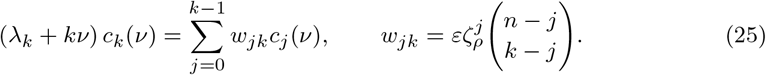

*Proof* On the simplex, write

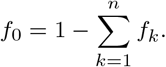

At *U* = 0, the Jacobian of the reduced vector field with respect to (*f*_1_, …, *f*_*n*_), evaluated at the pristine equilibrium, is upper bidiagonal. Its diagonal entries are

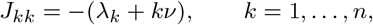

and its superdiagonal entries are

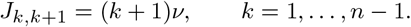

Since *λ*_*k*_+ *kν >* 0, every diagonal entry is strictly negative, so *J* is invertible. Since 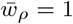 at the pristine state and remains positive in a neighborhood of it, the steady-state map is analytic in (*f, U*) near (*f, U*) = ((1, 0, …, 0), 0). The implicit function theorem therefore gives a locally unique equilibrium branch analytic in *U* .

Write this branch as

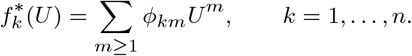

We show by induction on *m* that

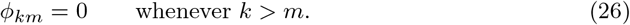

At order *U* ^*m*^, the steady-state equations have the form

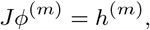

where *ϕ*^(*m*)^ = (*ϕ*_1*m*_, …, *ϕ*_*nm*_)^*T*^ and *h*^(*m*)^ depends only on coefficients of lower order in *U* and on the explicit *U* -dependence of the mutation kernel.

Assume inductively that *ϕ*_*jr*_= 0 whenever *j > r* for all *r < m*. For a mutation term feeding class *k* from a lower class *j < k*,

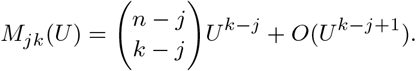

By the induction hypothesis, 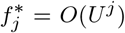 through order *m* − 1, so such a contribution has order at least

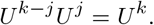

Hence no mutation inflow from below contributes to the coefficient of *U* ^*m*^ when *k > m*.

The branch is analytic in *U* and *f* ^∗^(0) is the pristine state, so 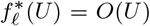 for every *l* ≥ 1 before any sharper hierarchy is imposed. Since 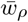and 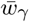 are smooth functions of *f* and both equal 1 at the pristine state, it follows that *R*^∗^ − *ε* = *O*(*U*). The corrections *R*^∗^ − *ε* and *M*_*kk*_(*U*) − 1 = *O*(*U*) in the same-class reproduction term multiply 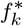, so their coefficient at order *U* ^*m*^ depends only on lower-order coefficients *ϕ*_*kr*_with *r < m*, which vanish when *k > m* by the induction hypothesis. Likewise, multiplying a mutation inflow from *j < k* by *R*^∗^ − *ε* can only raise its order beyond *U* ^*k*^ . Therefore

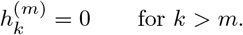

Because *J* is upper bidiagonal, backward substitution now gives

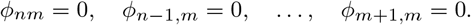

This proves (26). Consequently,

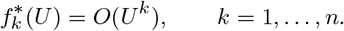

To obtain the leading coefficient, substitute 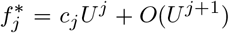 and *R*^∗^ = *ε* + *O*(*U*) into the class-*k* equilibrium equation. For *j < k*,

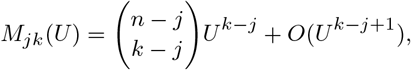

whereas the repair inflow 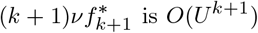. Collecting the terms of order *U* ^*k*^ gives (25). Because *c*_0_(*ν*) = 1, *w*_*jk*_*>* 0, and *λ*_*k*_+ *kν >* 0, the recursion (25) implies inductively that *c*_*k*_(*ν*) *>* 0 for every *k* 1. Hence, for sufficiently small *U >* 0, all 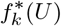are positive and 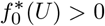, so the analytic branch lies in Δ_*n*_. □

#### Corollary 4

(Local stability of the rare-mutation branch) *For fixed n <* ∞ *and finite ν* ≥ 0, *there exists U*_0_ *>* 0 *such that the equilibrium branch of Proposition 3 is locally asymptotically stable within* Δ_*n*_*for*

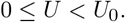

*Proof* At *U* = 0, the Jacobian of the reduced system is upper bidiagonal with eigenvalues

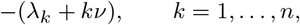

all of which are strictly negative. The pristine equilibrium is therefore a hyperbolic sink. Proposition 3 gives an equilibrium branch analytic in *U*, and the Jacobian evaluated along that branch depends continuously on *U* . Hence its eigenvalues remain in the open left half-plane for all sufficiently small *U* ≥ 0. □

From the exact no-repair equilibrium, (15), with *d* = *ζ*_*γ*_ − *ζ*_*ρ*_,

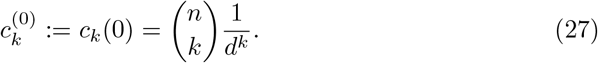

Thus the no-repair coefficients needed below are known explicitly rather than recursively.

### 4.2 Path-filtering theorem

Define the class-*k* repair filter

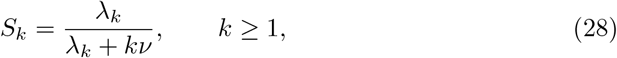

Here *λ*_*k*_ is the net demographic disadvantage of class *k* relative to the pristine state at the mutation-free homeostatic equilibrium. In particular,

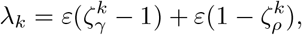

so it incorporates both excess mortality and reduced reproduction. It is therefore an effective demographic depletion rate, not a physical mortality hazard. The factor *S*_*k*_ measures the fraction of the leading-order class-*k* depletion coefficient attributable to this demographic disadvantage rather than repair. It should not be interpreted as the probability that an individual dies before being repaired.

Define the leading repair-suppression factors by

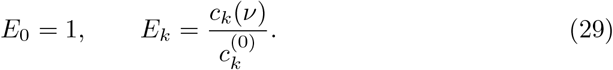

#### Theorem 1

(Repair-mediated path filtering) *For each fixed target class k* ∈ {1, …, *n*},

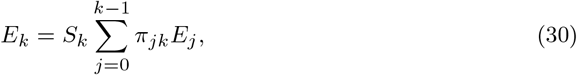

*Where*

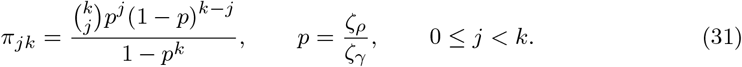

*The coefficients π*_*jk*_*are positive and sum to one. Equivalently, if J* ∼ Binomial(*k, p*), *then*

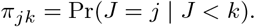

*Hence the conditional-binomial baseline weights π*_*jk*_*describing the leading no-repair source flux into class k are independent of the total number of loci n whenever n* ≥ *k. Since E*_0_ = 1 *and the recursion* (30) *depends on n only through these weights, the suppression factor E*_*k*_*is likewise independent of n*.

*Moreover*,

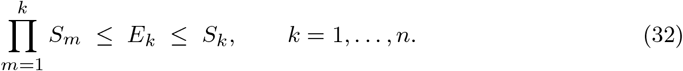

*If ν >* 0 *and k* ≥ 2, *both inequalities are strict:*

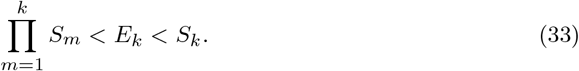

*For each fixed k* ≥ 1, *the suppression factor E*_*k*_(*ν*) *is strictly decreasing in ν* ≥ 0, *with E*_*k*_(0) = 1. *Finally, the class-k contribution to the equilibrium load satisfies*

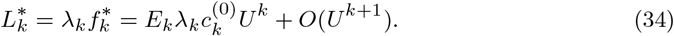

*Proof* From Proposition 3,

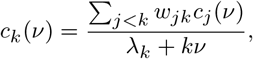

whereas at *ν* = 0,

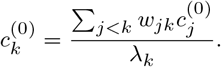

Dividing the two expressions and using 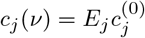gives

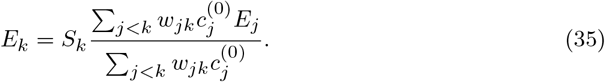

The exact no-repair coefficient (27) gives

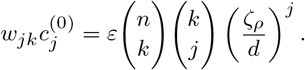

The factor 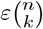cancels from (35). Since *d* = *ζ*_*γ*_− *ζ*_*ρ*_, normalization over *j < k* gives

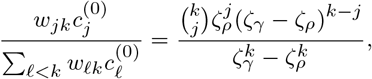

which is precisely (31) after writing *p* = *ζ*_*ρ*_*/ζ*_*γ*_. This proves (30) and also shows that the normalized baseline source weights, and therefore *E*_*k*_, are independent of *n* for every *n* ≥ *k*.

It remains to establish the bounds. For *k* = 1, only *j* = 0 contributes, so *E*_1_ = *S*_1_. Suppose inductively that

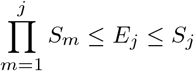

for every *j < k*. Since 0 *< S*_*j*_≤ 1, all terms in the weighted average in (30) satisfy *E*_*j*_≤ 1, and therefore

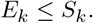

Similarly,

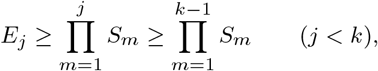

with the empty product equal to one for *j* = 0. Hence

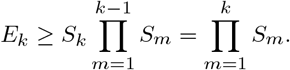

The induction closes, proving (32).

To establish monotonicity, observe that

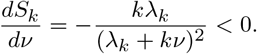

Since the weights *π*_*jk*_are positive and independent of *ν*, and *E*_0_ = 1, induction in (30) shows that *E*_*k*_(*ν*) is strictly decreasing for every *k* ≥ 1. At *ν* = 0, all *S*_*k*_= 1, so the same recursion gives *E*_*k*_(0) = 1.

Moreover, both inequalities are strict for *k* ≥ 2 and *ν >* 0. Since 0 *< p <* 1, every route weight *π*_*jk*_is positive. The weight *π*_0*k*_*>* 0 attaches to *E*_0_= 1, which strictly exceeds 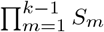 because *S*_*m*_*<* 1 for *ν >* 0. Hence

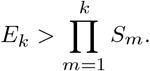

Similarly, *π*_1*k*_*>* 0 attaches to *E*_1_= *S*_1_*<* 1, while all *E*_*j*_≤ 1, so the weighted average in (30) is strictly less than one. Therefore

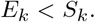

Thus both inequalities in (32) are strict for *k* ≥ 2 and *ν >* 0, which proves (33).

Finally, substituting 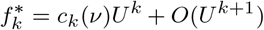 and 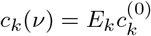 into 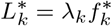 gives (34). □

*Remark* 1 (Refined bounds and the strong-repair limit). For *k ≥* 2, the recursion (30) and the bounds (32) yield

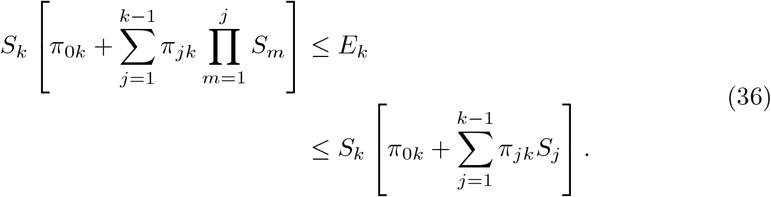

These bounds sharpen the original sandwich and coincide for *k* = 2.

Now let *ν* → ∞ with *n, k*, and the demographic parameters fixed. Since 0 *< E*_*j*_ ≤ *S*_*j*_ → 0 for every *j* ≥ 1, while *E*_0_ = 1, Eq. (30) gives

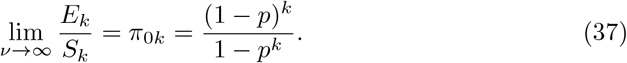

Using *p* = *ζ*_*ρ*_*/ζ*_*γ*_ and 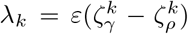, we obtain the more explicit asymptotic relation

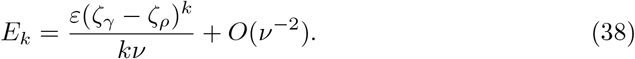

Thus, for *k* ≥ 2 and fixed 0 *< p <* 1, the terminal-filter bound remains separated from *E*_*k*_ by a nonzero relative gap in the strong-repair limit. The refined bounds in (36), however, are asymptotically sharp.

The leading surviving source contribution comes from direct 0 → *k* mutation events, which bypass intermediate repair filters. Consequently, *E*_*k*_ = *O*(*ν*^−1^), whereas the fully sequential lower bound 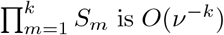For *k* = 1, the identity *E*_1_ = *S*_1_ continues to hold for every *ν*.

Define the leading-order no-repair contribution from class *k* by

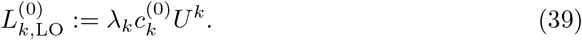

Then (34) can be written as

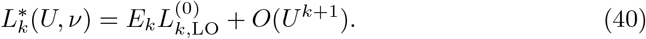

#### 4.2.1 Single-locus recovery and total leading load

For *k* = 1, Eq. (30) gives

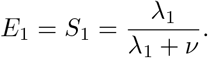

Since class 1 is the only *O*(*U*) contribution to the total load,

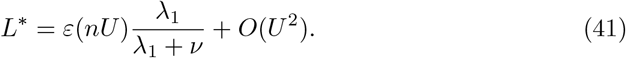

Thus the single-locus repair-modified Haldane result is recovered with the leading mutational input expressed through the genomic rate *nU* . This is also the scaling used in Corollary 3, although the expansion here is for fixed *n* and does not by itself establish a repaired genomic limit.

#### 4.2.2 Sequential and direct-route limits

Equation (31) makes the limiting source structures explicit. As *p* → 0, *π*_0*k*_ → 1, so

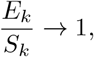

corresponding to concentration of the baseline source weight on direct 0 → *k* multilocus mutation.

As *p* → 1, *π*_*k*−1,*k*_ → 1, so the conditional baseline source law concentrates on the immediately preceding class, corresponding to sequential accumulation. For fixed *ν >* 0, this admissible limit also forces *ζ*_*γ*_ − *ζ*_*ρ*_ → 0, so *λ*_*j*_ → 0 and hence *S*_*j*_ → 0. Consequently, concentration of the source weigh∏ts on *j* = *k* − 1 should not be interpreted as the stronger relative statement 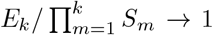; vanishing bypass weights can remain comparable to the already-small sequential contribution.

##### Corollary 5

(Repair-induced association among loci) *Let n* ≥ 2 *and ν >* 0. *Along the rare-mutation equilibrium branch, the leading pairwise association between distinct damage loci is positive. In particular, if X*_*l*_∈ {0, 1} *denotes the damage state of locus l, then for l*/= *m*,

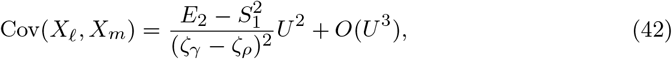

*Where*

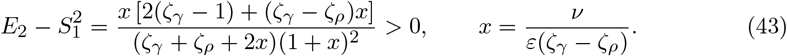

*Since* 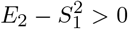, *it follows that* Cov(*X*_*l*_, *X*_*m*_) *>* 0 *for all sufficiently small U >* 0. *Hence, for sufficiently small U, the equilibrium damage number is overdispersed relative to a binomial distribution with the same marginal damage probability*.

*Proof* For *k* = 2, Eq. (30) gives

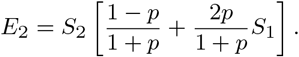

Writing

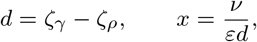

gives

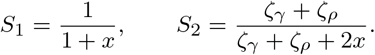

Direct simplification yields (43), which is strictly positive for *ν >* 0 because *ζ*_*γ*_*>* 1 and *ζ*_*γ*_*> ζ*_*ρ*_.

To justify the passage from class frequencies to labeled loci, we first establish exchange-ability of the local microstate equilibrium branch. Work on the microstate simplex with

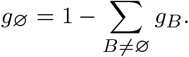

The microstate vector field is equivariant under permutations of the loci. At *U* = 0, the pristine state *g*_∅_= 1 is an equilibrium. Ordered by Hamming weight, the Jacobian in the coordinates (*g*_*B*_)_*B* ∅_is block upper bidiagonal: repair couples weight-*k* + 1 states only to weight-*k* states, while the diagonal block corresponding to |*B*| = *k* is

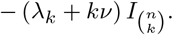

Since *λ*_*k*_+ *kν >* 0, this Jacobian is invertible. The implicit function theorem therefore gives a locally unique microstate equilibrium branch *g*^∗^(*U*) analytic near *U* = 0.

If *σ* ∈ S_*n*_is any permutation of the loci, equivariance implies that *σ* · *g*^∗^(*U*) is also an equilibrium branch near the pristine state. Local uniqueness therefore forces

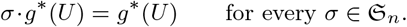

Thus the repaired equilibrium branch is exchangeable. Aggregating this microstate branch by Hamming weight produces an equilibrium branch of the class system, and local uniqueness in Proposition 3 implies that its class frequencies are precisely the 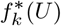 constructed there.

By exchangeability,

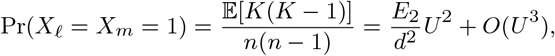

Whereas

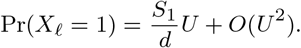

Subtracting the product of the marginals gives (42).

Writing

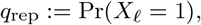

exchangeability gives

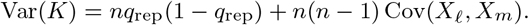

The positive covariance therefore implies overdispersion relative to the matched binomial distribution for sufficiently small *U* . □

### 4.3 Numerical validation

To compare the finite-*U* nonlinear equilibrium with the leading-order path-filtering prediction, define the exact no-repair class contribution at the same mutation probability by

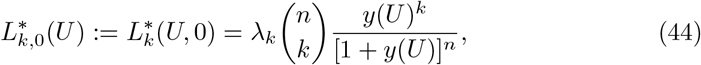

and, for *U >* 0, the normalized repair-suppression ratio

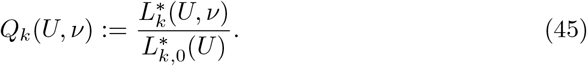

Since

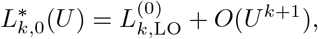

Eq. (40) implies

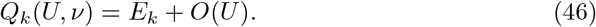

Thus the finite-*U* suppression ratio converges to the path-filtering factor *E*_*k*_ with an *O*(*U*) error.

For reproducibility, the numerical validation used *n* = 4, *ε* = 0.5, *ζ*_*ρ*_ = 0.8, *U* = 0.01, *ζ*_*γ*_ ∈ { 1.15, 1.30, 1.50}, and *ν* ∈ [0, 1] sampled at 41 equally spaced values. Direct integrations of (8) were performed in R (R Core Team 2024) with the LSODA implementation in the <monospace>deSolve</monospace> package (Soetaert et al. 2010) from the pristine initial condition

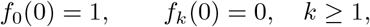

to terminal time *τ*_max_ = 20000, with relative tolerance 10^−10^ and absolute tolerance 10^−12^. Steady state was assessed by evaluating the right-hand side of the ODE system at the terminal state and requiring

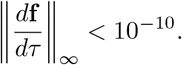

For these parameters, the finite-*U* ratios *Q*_*k*_(*U, ν*) closely follow the leading-order predictions *E*_*k*_ over the full range *ν* ∈ [0, 1] and across the three fitness landscapes *ζ*_*γ*_ ∈ { 1.15, 1.30, 1.50} (Fig. 1). The separation among landscapes reflects the dependence of the repair-suppression factors on the demographic fitness parameters.

**Fig. 1.**
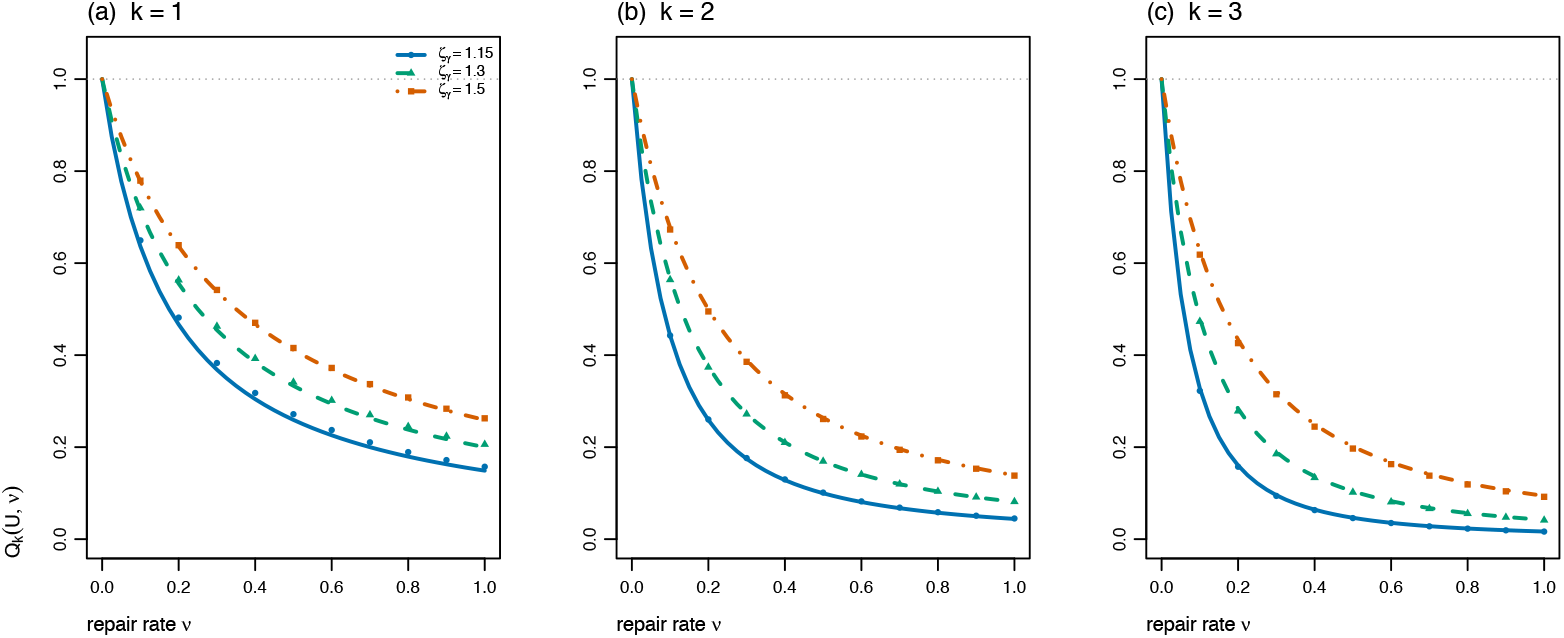
Numerical validation of repair-mediated path filtering for the full nonlinear *n* = 4 system with *ε* = 0.5, *ζ*_*ρ*_ = 0.8, and *U* = 0.01. Panels show the finite-*U* suppression ratio 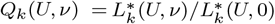 as a function of repair rate *ν* for *k* = 1 (left), *k* = 2 (center), and *k* = 3 (right). Lines show the leading-order predictions *E*_*k*_ for *ζ*_*γ*_ ∈ { 1.15, 1.30, 1.50}, distinguished by solid blue, dashed green, and dot-dashed orange lines, respectively. Filled circles, triangles, and squares show the corresponding finite-*U* ratios obtained by direct LSODA integration of the full class-frequency system, with the exact no-repair equilibrium used in the denominator. The light-gray dotted line marks the unrepaired value *Q*_*k*_ = 1. Agreement between the numerical ratios and analytical predictions is consistent with *Q*_*k*_(*U, ν*) = *E*_*k*_ + *O*(*U*).

To test the predicted remainder directly, we fixed the representative parameter slice *ζ*_*γ*_ = 1.50 and *ν* = 0.5, repeated the calculation at

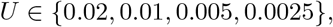

and evaluated

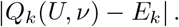

The residual decreased with slope approximately one on a log–log scale for *k* = 1, 2, 3, consistent with the *O*(*U*) remainder in (46) (Online Resource 1, Supplementary Fig. S1).

As an additional check of the *n*-independence predicted by Theorem 1, we fixed the target class *k* = 3 and repeated the nonlinear integration for *n* ∈ {3, 4, 6, 8} at *U* = 0.01. The resulting finite-*U* ratios lie close to the same analytical curve *E*_3_(*ν*) (Online Resource 1, Supplementary Fig. S2), consistent with the theorem’s prediction that the leading suppression factor for fixed *k* is independent of *n*.

## 5 Discussion

The multilocus model retains considerably more exact structure than might be expected from its nonlinear homeostatic feedback. Permutation symmetry reduces the 2^*n*^ microstate dynamics exactly to *n* + 1 damage classes. In the absence of repair, the nonlinear class-equilibrium set admits a complete classification, including explicit shifted-binomial distributions on invariant boundary faces. On the pristine-positive branch, the corresponding microstate equilibrium is an exact Bernoulli product measure, with a binomial damage number and a Poisson limit under genomic rare-mutation scaling. Repair changes this structure qualitatively: it breaks the no-repair product factorization and introduces route-dependent filtering of mutation load.

The path-filtering theorem identifies the mechanism by which repair modifies mutation–selection balance across multiple damage levels. Unlike a purely sequential hierarchy, the multilocus system permits direct transitions from a source class *j* to a target class *k > j* + 1, allowing mutation to bypass intermediate classes and hence some of their repair filters. Consequently, for *k* ≥ 2 and *ν >* 0, the leading suppression factor *E*_*k*_ lies strictly between the fully sequential product 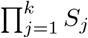 and the terminal filter *S*_*k*_. The conditional-binomial source weights *π*_*jk*_ quantify how the realized suppression interpolates between these limiting route structures.

As *ν* → ∞, each *S*_*j*_ → 0, and hence *E*_*k*_ → 0 for every fixed damage class. The ordering of the suppression factors is parameter-dependent rather than universal. For example, at *ε* = 0.5, *ζ*_*ρ*_ = 0.8, *ζ*_*γ*_ = 2.0, and *ν* = 0.5, Eq. (30) gives

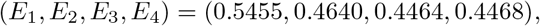

so *E*_4_ *> E*_3_. With *ζ*_*γ*_ = 3.0 and *ν* = 0.1, the sequence becomes

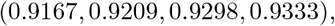

which is strictly increasing.

A steeper demographic landscape increases the net disadvantage *λ*_*k*_ relative to the repair coefficient *kν*, while also shifting the conditional source weights toward direct multilocus jumps. Both effects can weaken the relative suppression of higher damage classes.

### 5.1 Repair generates statistical association among loci

Without repair, the equilibrium microstate distribution is exactly a Bernoulli product measure (Corollary 2), so distinct loci are statistically independent. Repair changes this qualitatively. As shown in Corollary 5, for every *ν >* 0 the leading pairwise covariance is positive:

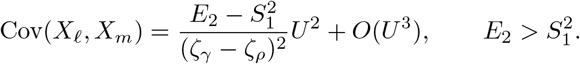

Thus, along the rare-mutation branch, repair breaks the exact no-repair factorization and produces an overdispersed damage distribution for sufficiently small *U >* 0.

The contrast with Hill–Robertson interference (Hill and Robertson 1966) is therefore one of mechanism rather than analogy. In the present deterministic homeostatic model, mutation and selection alone yield independent loci at the pristine-positive equilibrium, despite their common demographic environment. Adding repair destroys this equilibrium product structure and generates positive leading-order association among loci.

### 5.1 Repair beyond the no-repair threshold

Proposition 2 shows that without repair, *U ≥ U*_*c*_ leaves the fully damaged state *e*_*n*_ as the only equilibrium. Positive repair changes this conclusion: every equilibrium for *ν >* 0 must satisfy 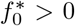. Indeed, if 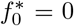, the class-0 balance forces 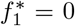, and successive class balances force all frequencies to vanish, contradicting normalization.

Numerically, for *n* = 4, *ε* = 0.5, *ζ*_*ρ*_ = 0.8, and *ζ*_*γ*_ = 1.3, so that *U*_*c*_ = 0.3846, we obtain 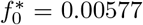 and 0.18626 at *U* = 0.5 for *ν* = 0.05 and 0.3, respectively. Even at *U* = 0.9, the corresponding frequencies remain positive: 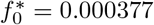 and 0.07050.

Thus positive repair prevents complete loss of the pristine class at equilibrium, even beyond the no-repair threshold. This does not, by itself, establish global attraction of the repaired equilibrium.

### 5.3 Multi-hit carcinogenesis

The suppression structure connects naturally to multi-hit models of carcinogenesis (Knudson 1971; Armitage and Doll 1954; Beerenwinkel et al. 2007; Foo et al. 2014; Portier and Kopp-Schneider 1991). In a homeostatic somatic tissue, the leading rare-mutation frequency of cells carrying *k* repairable damage states is suppressed relative to its no-repair value by the factor *E*_*k*_, which reflects both the depletion of intermediate reservoirs by repair and the partial bypass of those reservoirs by simultaneous multilocus mutations. The sandwich ∏*S*_*j*_ ≤ *E*_*k*_ ≤ *S*_*k*_ therefore has a concrete leading-order implication: the repair-suppression factor for the *k*-hit frequency lies between the fully sequential prediction (smaller *E*_*k*_, stronger suppression) and the terminal-filter prediction (larger *E*_*k*_, weaker suppression). The realized suppression *E*_*k*_ interpolates between these limits according to the baseline mixture of source classes encoded by *π*_*jk*_. The conditional-binomial source law makes this interpolation explicit. At leading order, the relative weight of sequential and direct routes is governed by *p* = *ζ*_*ρ*_*/ζ*_*γ*_: as *p* → 1, source weight concentrates near *j* = *k* − 1, whereas as *p* → 0, source weight concentrates near *j* = 0 and the terminal-filter bound is approached. Thus the mutation probability sets the abundance of rare *k*-hit states, while the demographic fitness contrast controls the leading mixture of routes by which they are reached.

In applications to multi-hit carcinogenesis, the damage states should be interpreted as reversible precursor states or effective molecular lesions, rather than necessarily as fixed driver mutations. The path-filtering result therefore characterizes the accumulation and restoration of repairable damage states that may precede irreversible genetic alterations. An explicit model of mutation fixation would require additional states distinguishing repairable lesions from established sequence changes.

### 5.4 Scope and extensions

The present analysis assumes identical loci, multiplicative demographic effects across damage loci, and a uniform per-locus repair rate *ν*. Heterogeneity among loci is known to alter mutation–selection equilibria in other multilocus settings (Waxman and Welch 2003). These symmetries allow the 2^*n*^ microstate system to close exactly on the *n* + 1 Hamming-weight classes and lead to the path-filtering recursion (30). If mutation, repair, reproduction, or mortality effects vary across loci, total damage number will generally no longer provide a closed state description. The same basic competition between repair filtering and mutational bypass may persist, but the relevant paths would then have to be resolved on the labeled microstate space rather than through a single damage-class recursion. Likewise, nonmultiplicative fitness effects would alter both the source weights and the class-specific filters and could generate qualitatively different route structure (Charlesworth 1990; Hermisson et al. 2002).

The rare-mutation expansion above treats *ν* as fixed as *U* → 0. A related regime arises when repair scales with mutation, *ν* = *θU*, for fixed *θ* ≥ 0. In this case, the repair outflux from class *k* enters at order *U*^*k*+1^ rather than at the leading order *U*^*k*^. Consequently, for fixed *n*, the leading multilocus load satisfies

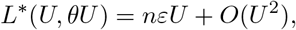

consistent with the corresponding result *L*^∗^ = *εU* + *O*(*U* ^2^) established in the one-dimensional hierarchy (Hardisty 2026). The first repair-dependent corrections therefore enter at higher order and require a joint expansion in mutation and repair to characterize explicitly. Physiological costs of reducing mutation have been discussed in the mutation-rate evolution literature (Sniegowski et al. 2000); here such costs are deliberately excluded from the state-restoration process. Incorporating such costs directly into reproduction or mortality would be a natural extension, but would change the homeostatic feedback and should be treated as part of the demographic model rather than appended to the present load calculation.

The role of locus number is more structured than might be expected. In the no-repair model, the genomic rare-mutation limit *n* → *∞, U* →0, with *nU* → Θ produces the Poisson equilibrium of Corollary 3. With repair, Theorem 1 shows that for every fixed target class *k*, the conditional baseline source weights and the leading suppression factor *E*_*k*_ are already independent of *n* for all *n* ≥ *k*. Thus increasing the total number of available loci changes the abundance of a fixed damage class through the factor 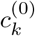 but not its leading repair-suppression factor.

The genomic rare-mutation limit requires particular care because the fixed-*n* rare-mutation expansion is not uniform in genome size. Indeed, even without repair,

Corollary 2 gives

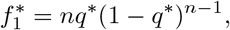

and therefore, under *U* = Θ*/n*,

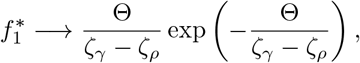

rather than the expression Θ*/*(*ζ*_*γ*_ − *ζ*_*ρ*_) obtained by retaining only the leading fixed-*n* term.

With repair, the same issue prevents the *n*-independence of *E*_*k*_ from establishing a genomic equilibrium limit. For example, at *ε* = 0.5, *ζ*_*ρ*_ = 0.8, *ζ*_*γ*_ = 1.3, *ν* = 0.4, and *n* = 32, the ratio of the numerically computed 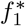 to *E*_1_Θ*/*(*ζ*_*γ*_ *™ ζ*_*ρ*_) is 0.882, 0.959, and 0.987 for Θ = 0.12, 0.04, and 0.012, respectively.

These observations show why the fixed-*n* path-filtering expansion does not by itself determine the repaired genomic limit. Establishing the existence and form of that limiting distribution requires a separate analysis.

Finally, the present results concern equilibrium under a constant mutation probability and repair rate. The local stability result established above is confined to the rare-mutation branch near *U* = 0; global attraction and stability farther from *U* = 0 remain open. Extending the path-filtering framework to time-dependent mutation or repair, or to damage accumulation over a finite tissue lifespan, would connect the equilibrium theory developed here to transient exposure histories and age-dependent multi-hit processes. Temporal variation in mutation itself has recently been considered in mutation-rate modifier models (Heinrich-Mora and Feldman 2026), but the corresponding problem for repair-mediated state restoration remains open.

## Acknowledgements

The author thanks Fred Adler and Jon Seger for many helpful discussions on population genetics and mathematical modeling.

## Statements and Declarations

### Funding

No funds, grants, or other support were received for conducting this study.

### Competing interests

The author has no relevant financial or non-financial interests to disclose.

### Data availability

No external datasets were used in this study. Numerical results reported in this article were generated from the mathematical model described in the manuscript.

### Code availability

The R code used for numerical integration and figure generation is archived on Zenodo at https://doi.org/10.5281/zenodo.23292998.

### Author contributions

The author conceived the study, developed the mathematical model, performed the analytical and numerical work, and wrote the manuscript.

### Ethics approval

Not applicable.

### Consent to participate

Not applicable.

### Consent for publication

Not applicable.

**Materials availability**

Not applicable.

### Use of generative AI and AI-assisted technologies

During the preparation of this work, the author used Claude Opus (Anthropic) and ChatGPT (OpenAI) to assist with manuscript editing, language refinement, organizational suggestions, and discussion of mathematical exposition. The scientific ideas, mathematical derivations, model development, and conclusions are the author’s own. The author reviewed and edited all AI-assisted content and takes full responsibility for the content of the manuscript.

## Footnotes

1 Here = 1 (1 *U*)^*n*^ is the mutation-escape probability of the *pristine* genome, not the population-average probability of an upward mutation event. The realized upward mutational flux is 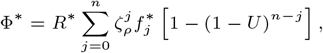 which is strictly smaller than 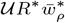 for 0 *< U < U*_*c*_, because damaged parents have fewer undamaged loci available to mutate. Thus (16) follows from the pristine-class balance that fixes *R*^∗^; it should not be interpreted as an equality between load and the realized upward mutational flux.

## Notes

https://github.com/Priceeqn1/Rdiffeqnscripts

### Competing Interest Statement

The authors have declared no competing interest.

## References

Armitage, P., Doll, R.: The age distribution of cancer and a multi-stage theory of carcinogenesis. British Journal of Cancer 8, 1–12 (1954) 10.1038/bjc.1954.1

Beerenwinkel, N., Antal, T., Dingli, D., Traulsen, A., Kinzler, K.W., Velculescu, V.E., Vogelstein, B., Nowak, M.A.: Genetic progression and the waiting time to cancer. PLoS Computational Biology 3, 225 (2007) 10.1371/journal.pcbi.0030225

Bürger, R., Hofbauer, J.: Mutation load and mutation–selection balance in quantitative genetic traits. Journal of Mathematical Biology 32, 193–218 (1994) 10.1007/BF00163878

Baake, E., Wagner, H.: Mutation–selection models solved exactly with methods of statistical mechanics. Genetical Research 78, 93–117 (2001) 10.1017/S0016672301005110

Charlesworth, B.: Mutation–selection balance and the evolutionary advantage of sex and recombination. Genetical Research 55, 199–221 (1990) 10.1017/S0016672300025532

Crow, J.F.: Genetic loads and the cost of natural selection. In: Kojima, K.-I. (ed.) Mathematical Topics in Population Genetics, pp. 128–177. Springer, Berlin (1970). 10.1007/978-3-642-46244-3_5

Chatterjee, N., Walker, G.C.: Mechanisms of DNA damage, repair, and mutagenesis. Environmental and Molecular Mutagenesis 58, 235–263 (2017) 10.1002/em.22087

Foo, J., Leder, K., Ryser, M.D.: Multifocality and recurrence risk: A quantitative model of field cancerization. Journal of Theoretical Biology 355, 170–184 (2014) 10.1016/j.jtbi.2014.02.042

Haigh, J.: The accumulation of deleterious genes in a population—muller’s ratchet. Theoretical Population Biology 14, 251–267 (1978) 10.1016/0040-5809(78)90027-8

Haldane, J.B.S.: The effect of variation of fitness. The American Naturalist 71, 337–349 (1937) 10.1086/280722

Hardisty, B.: Homeostatic Turnover, Repair, and the Demographic Form of Haldane’s Principle. bioRxiv preprint (2026). 10.64898/2026.09.20.753009

Hochedlinger, K., Jaenisch, R.: Induced pluripotency and epigenetic reprogramming. Cold Spring Harbor Perspectives in Biology 7, 019448 (2015) 10.1101/cshperspect.a019448

Heinrich-Mora, E., Feldman, M.W.: Evolution under stochastic transmission: Mutation-rate modifiers. Theoretical Population Biology 168, 65–78 (2026) 10.1016/j.tpb.2026.02.003

Hill, W.G., Robertson, A.: The effect of linkage on limits to artificial selection. Genetical Research 8, 269–294 (1966) 10.1017/S0016672300010156

Hermisson, J., Redner, O., Wagner, H., Baake, E.: Mutation–selection balance: Ancestry, load, and maximum principle. Theoretical Population Biology 62, 9–46 (2002) 10.1006/tpbi.2002.1582

Johnson, T.: The approach to mutation–selection balance in an infinite asexual population, and the evolution of mutation rates. Proceedings of the Royal Society of London B 266, 2389–2397 (1999) 10.1098/rspb.1999.0936

Kimura, M., Maruyama, T.: The mutational load with epistatic gene interactions in fitness. Genetics 54, 1337–1351 (1966) 10.1093/genetics/54.6.1337

Knudson, A.G.: Mutation and cancer: Statistical study of retinoblastoma. Proceedings of the National Academy of Sciences of the United States of America 68, 820–823 (1971) 10.1073/pnas.68.4.820

Lindahl, T.: Instability and decay of the primary structure of DNA. Nature 362, 709–715 (1993) 10.1038/362709a0

Portier, C.J., Kopp-Schneider, A.: A multistage model of carcinogenesis incorporating DNA damage and repair. Risk Analysis 11, 535–543 (1991) 10.1111/j.1539-6924.1991.tb00639.x

R Core Team: R: A Language and Environment for Statistical Computing. R Foundation for Statistical Computing, Vienna, Austria (2024). https://www.R-project.org/

Sniegowski, P.D., Gerrish, P.J., Johnson, T., Shaver, A.: The evolution of mutation rates: Separating causes from consequences. BioEssays 22, 1057–1066 (2000) 10.1002/1521-1878(200012)22:12‹1057::AID-BIES3›3.0.CO;2-W

Soetaert, K., Petzoldt, T., Setzer, R.W.: Solving differential equations in r: Package desolve. Journal of Statistical Software 33(9), 1–25 (2010) 10.18637/jss.v033.i09

Waxman, D., Welch, J.: Non-equivalent loci and mutation–selection balance. Theoretical Population Biology 63, 339–345 (2003) 10.1016/S0040-5809(03)00011-X

