## Supplementary Figures for "Repair-Mediated Path Filtering of Mutation Load in Multilocus Homeostatic Populations"

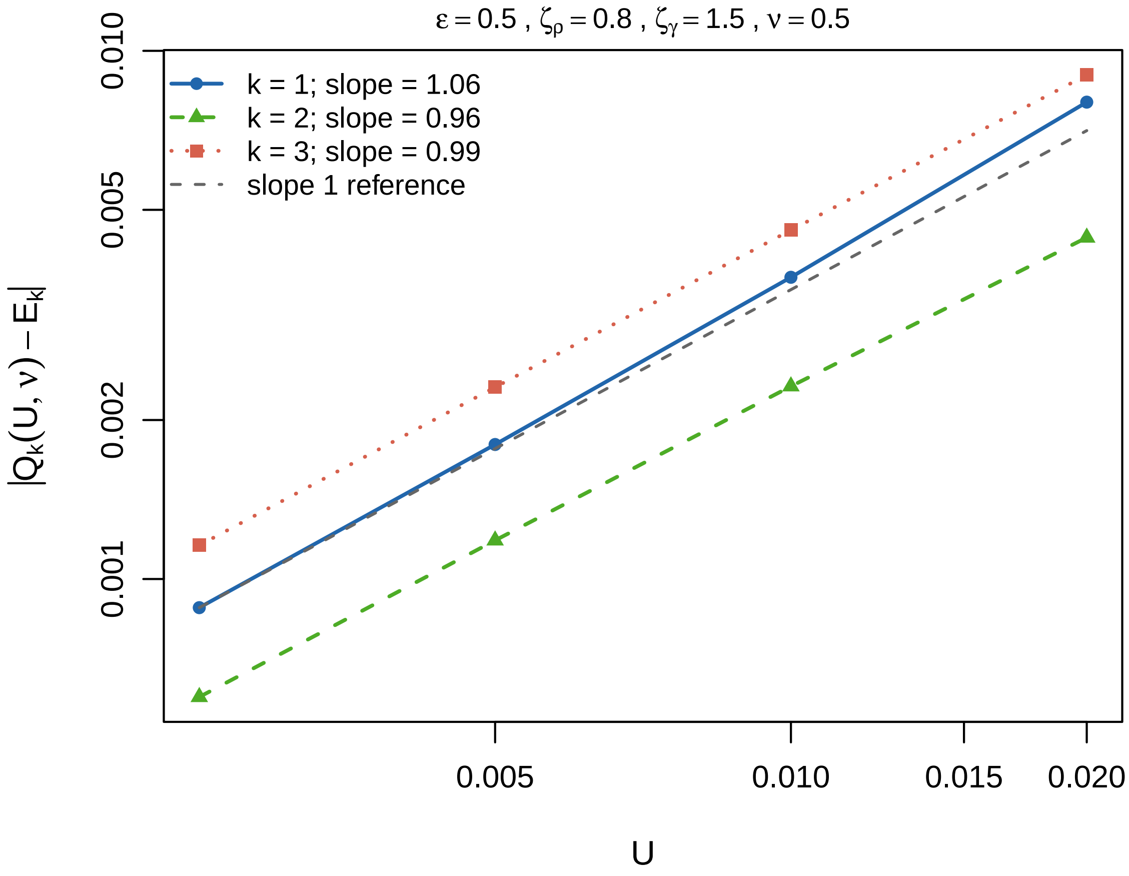


**Fig. S1** Convergence of the leading-order path-filtering prediction with mutation probability. For a representative slice ($\varepsilon = 0.5$, $\zeta_\rho = 0.8$, $\zeta_\gamma = 1.50$, $\nu = 0.5$), the residual $|Q_k(U,\nu) - E_k|$ is plotted against $U$ on log–log axes for $k = 1, 2, 3$, with $Q_k(U,\nu)$ obtained from the full nonlinear $n = 4$ steady state and the exact no-repair equilibrium used in the denominator. Each residual decreases approximately linearly with $U$, with fitted log–log slopes of approximately 1.06, 0.96, and 0.99 for $k = 1, 2, 3$, respectively, consistent with the prediction $Q_k(U,\nu) = E_k + O(U)$ in the main manuscript.

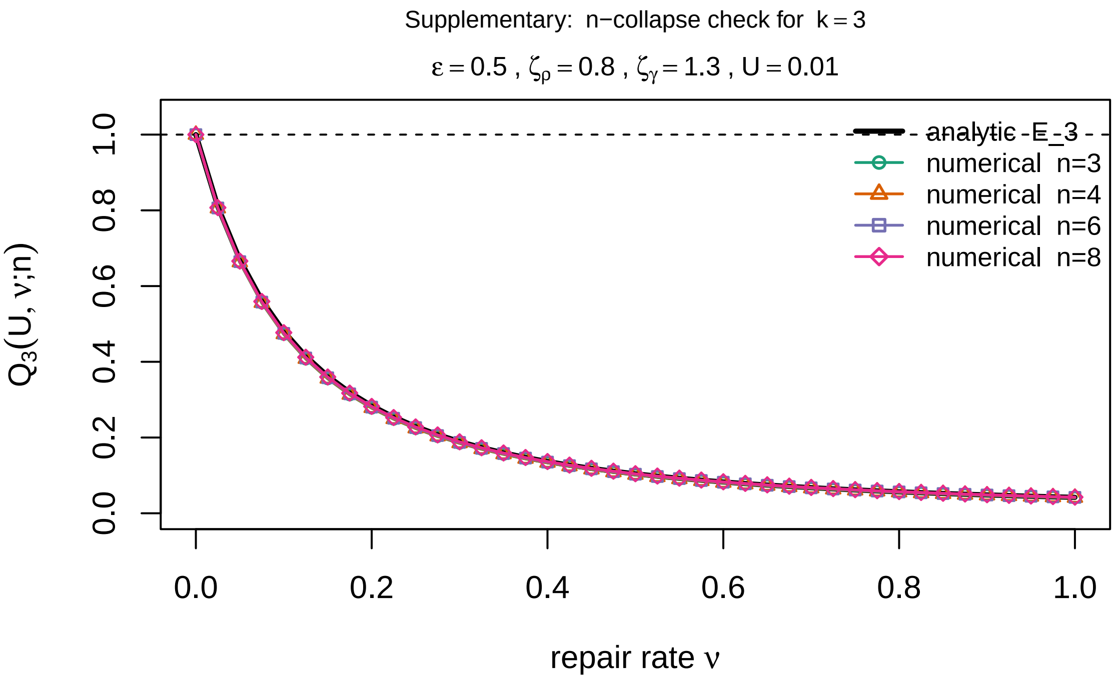


**Fig. S2** Supplementary numerical check of the $n$-independence predicted by Theorem 1 of the main manuscript for fixed target class $k = 3$. The solid black curve shows the leading-order path-filtering prediction $E_3(\nu)$, which is independent of the total number of loci $n$ for all $n \geq 3$. Colored curves and symbols show the finite-$U$ nonlinear suppression ratio $Q_3(U, \nu; n) = L_3^*(U, \nu; n)/L_3^*(U, 0; n)$, obtained by direct integration of the full class-frequency system for $n \in \{3, 4, 6, 8\}$, with $U = 0.01$, $\varepsilon = 0.5$, $\zeta_\rho = 0.8$, and $\zeta_\gamma = 1.30$. All four finite-$U$ curves lie close to the same analytical prediction, consistent with $Q_3(U, \nu; n) = E_3(\nu) + O(U)$ and with the theorem's result that the leading repair-suppression factor for a fixed damage class is independent of the total locus number.
